# Surveillance for SARS-CoV-2 in Ohio’s wildlife, companion, and agricultural animals

**DOI:** 10.1101/2022.12.30.522311

**Authors:** Margot Ehrlich, Christopher Madden, Dillon S. McBride, Jaqueline M. Nolting, Devra Huey, Scott Kenney, Qiuhong Wang, Linda Saif, Anastasia Vlasova, Patricia Dennis, Dusty Lombardi, Stormy Gibson, Alexis McLaine, Sarah Lauterbach, Page Yaxley, Jenessa Winston, Dubraska Diaz-Campos, Risa Pesapane, Mark Flint, Jaylene Flint, Randy Junge, Seth A. Faith, Andrew S. Bowman, Vanessa L. Hale

## Abstract

Severe acute respiratory syndrome coronavirus 2 (SARS-CoV-2) emerged in humans in late 2019 and spread rapidly to become a global pandemic. A zoonotic spillover event from animal to human was identified as the presumed origin. Subsequently, reports began emerging regarding spillback events resulting in SARS-CoV-2 infections in multiple animal species. These events highlighted critical links between animal and human health while also raising concerns about the development of new reservoir hosts and potential viral mutations that could alter virulence and transmission or evade immune responses. Characterizing susceptibility, prevalence, and transmission between animal species became a priority to help protect animal and human health. In this study, we coalesced a large team of investigators and community partners to surveil for SARS-CoV-2 in domestic and free-ranging animals around Ohio between May 2020 and August 2021. We focused on species with known or predicted susceptibility to SARS-CoV-2 infection, highly congregated or medically compromised animals (e.g. shelters, barns, veterinary hospitals), and animals that had frequent contact with humans (e.g. pets, agricultural animals, zoo animals, or animals in wildlife hospitals). This included free-ranging deer (n=76), mink (n=57), multiple species of bats (n=65), and other wildlife in addition to domestic cats (n=275) and pigs (n= 184). In total, we tested 800 animals (34 species) via rRT-PCR for SARS-CoV-2 RNA. SARS-CoV-2 viral RNA was not detected in any of the tested animals despite a major peak in human SARS-CoV-2 cases that occurred in Ohio subsequent to the peak of animal samplings. Importantly, due to lack of validated tests for animals, we did not test for SARS-CoV-2 antibodies in this study, which limited our ability to assess exposure. While the results of this study were negative, the surveillance effort was critical and remains key to understanding, predicting, and preventing re-emergence of SARS-CoV-2 in humans or animals.

## Main text

As of December 2022, there have been over 649 million confirmed cases of COVID-19 worldwide [1]. SARS-CoV-2, the virus which causes COVID-19, infects a broad range of hosts species (n=25) [2]. Susceptibility to and transmission of SARS-CoV-2 within and between a growing list of species raises concerns regarding the development of new reservoir hosts, the re-emergence of COVID-19 from these species, and the increased potential for viral mutations that evade immune responses. Experimental animal work and ACE2 models also show the potential for broad host-susceptibility [3,4]. Early in the pandemic, along with rising SARS-CoV-2 prevalence in humans, our goal was to assess prevalence and transmission risk across a wide range of animal species, both under human care and free-ranging. We focused on species with known or predicted susceptibility to SARS-CoV-2; animals in high-risk situations (e.g., densely congregated, medically compromised); and animals that had frequent contact with humans or human environments - pets, agricultural animals, wildlife, animals in zoos or hospitals. A total of 800 animals were sampled from May 2020–August 2021. Sampling locations included: Ohio Wildlife Center, Columbus Zoo & Aquarium, Ohio State University Veterinary Medical Center, MedVet Hilliard, Columbus Humane, Shelter Outreach Services of Ohio, and Ohio county fairs and metroparks. Private citizens—hunters / trappers—also collected samples for this study.

Depending on species and sampling conditions, nasal, oropharyngeal, choanal, conjunctival, and/or rectal swabs (Fisherbrand^™^ Synthetic-Tipped Applicators) were collected from each animal. For species identified as high risk for SARS-CoV-2 (e.g., cats, ferrets), oropharyngeal, conjunctival, nasal, and rectal swabs were collected, as feasible; for pigs and deer - nasal swabs. Swabs were placed in brain heart infusion broth, viral transport media, or RNAlater^™^ and frozen at -80°C where they remained until extraction. When freezers were not immediately available, samples were placed on ice for up to 12 hours or into liquid nitrogen. A few animals (n=8) were tested on multiple dates, and counted as separate individuals as infection could have occurred between testing dates. SARS-CoV-2 rRT-PCR testing was conducted as described previously [5].

We sampled 34 species over 16 months, and SARS-CoV-2 viral RNA was not detected in any sample (**Table 1**). Human infection rates in Ohio varied widely during this time, peaking in December 2020 at 255,965 cases (2.17% prevalence) [6]. Despite the human caseload of SARS-CoV-2 and although many of the species tested demonstrated moderate to high susceptibility to SARS-CoV-2, none of the 800 animals tested positive.

**Table 1.** Species in Ohio tested for SARS-CoV-2 (May 2020-August 2021).

| Scientific Name | Common Name | Number<br>Individuals<br>Tested | Provenance<br>(HC = under human care;<br>FR = free-ranging) |
| --- | --- | --- | --- |
| *Acinonyx jubatus* | Cheetah | 2 | HC |
| *Branta canadensis* | Canada Goose | 1 | HC |
| *Canis latrans* | Coyote | 1 | FR |
| *Castor canadensis* | American Beaver | 4 | FR |
| *Colobus guereza* | Guereza Colobus<br>Monkey | 1 | HC |
| *Didelphis virginiana* | Virginia<br>Opossum | 7 | HC |
| *Eptesicus fuscus* | Big Brown Bat | 61 | HC |
| *Felis catus* | Domestic Cat* | 275 | pets: n=84;<br>shelter: n=191 |
| *Hydrochoerus hydrochaeris* | Capybara | 1 | HC |
| *Lasionycteris noctivagans* | Silver-haired bat | 1 | HC |
| *Lasiurus borealis* | Eastern Red Bat | 2 | HC |
| *Lemur catta* | Ringtailed lemur | 1 | HC |
| *Lontra canadensis* | North American<br>River Otter | 1 | HC |
| *Mandrillus sphinx* | Mandrill | 1 | HC |
| *Marmota monax* | Groundhog | 3 | HC |
| *Mephitis mephitis* | Striped Skunk | 25 | HC |
| *Mustela furo* | Domestic Ferret | 6 | pets |
| *Neogale vison* | American Mink | 58 | FR: n=57; HC: n=1 |
| *Odocoileus virginianus* | White-tailed<br>deer** | 76 | FR |
| *Ondatra zibethicus* | Common Muskrat | 36 | FR: n=57; HC: n=1 |
| *Pan paniscus* | Bonobo | 1 | HC |
| *Perimyotis subflavus* | Tricolored Bat | 1 | HC |
| *Pongo pygmaeus* | Bornean Orangutan | 1 | HC |
| *Potos flavus* | Kinkajou | 1 | HC |
| *Procyon lotor* | Raccoon | 15 | FR: n=57; HC: n=1 |
| *Sciurus carolinensis* | Eastern Grey Squirrel | 11 | HC |
| *Sciurus niger* | Eastern Fox Squirrel | 6 | HC |
| *Sus scrofa domesticus* | Domestic Pig | 184 | agricultural |
| *Sylvilagus floridanus* | Eastern Cottontail Rabbit | 12 | HC |
| *Tamias striatus* | Eastern Chipmunk | 1 | HC |
| *Tamiasciurus hudsonicus* | American Red Squirrel | 1 | HC |
| *Trachypithecus cristatus* | Silvered Leaf Langur | 1 | HC |
| *Turdus migratorius* | American Robin | 1 | HC |
| *Varecia rubra* | Red ruffed lemur | 1 | HC |

\*A subset of these cats are described in more detail (J.Winston, in preparation)
\*\*Does not include Ohio deer described in [5].

## Discussion

Although case reports of animal infections emerged throughout the pandemic, our early active surveillance efforts did not yield any SARS-CoV-2 detections. While it is possible that we missed viral shedding windows, infection dynamics can also change over time. For example, the 78 free-ranging deer tested in this study were sampled October-November 2020 and all were negative. In a related study conducted January-March 2021, 35% (129 of 360) of the free-ranging Ohio deer tested positive for SARS-CoV-2 [5]. The 2021 study occurred after the human COVID-19 peak and after deer gun hunting season in Ohio, both of which would have created additional opportunities for direct and indirect contact (e.g., environmental contamination) between humans and deer. Additional reports nationwide confirmed widespread infection in deer [7,8]. Notably, it took time to detect natural infections in deer. Negative results in early surveillance efforts did not preclude a potential public health threat associated with animal reservoir establishment. In fact, the threat of re-emergence from animal hosts has already been realized through mink-, hamster-, cat-, and probable deer-to-human transmission events [9-13]. These cases highlight the critical need for continued surveillance.

Mink, bats, and deer have been identified as potential reservoirs for SARS-CoV-2. However, most SARS-CoV-2 sampling efforts have focused on farmed rather than free-ranging mink [13]. Assessing viral prevalence in free-ranging animals is critical to determining the SARS-CoV-2 reservoir and transmission potential of species like mink. This Ohio study is the largest sampling effort, to date, in free-ranging mink (n=57) and in native bats (n=65, 4 species).

Deer mice are widespread across Ohio and North America and also have the potential to act a host for SARS-CoV-2 [14]. Limited surveillance in these free-ranging species could mean undetected virus circulation and maintenance in the environment. Although our study included 105 samples from other rodent species, no samples from deer mice were obtained, highlighting key gaps in our surveillance efforts. Targeted efforts to surveil free-ranging deer mice will be necessary to evaluate infection and transmission in these species as well as their potential to serve as an intermediate host for other animals like white-tailed deer. One limitation of this study: We only tested for SARS-CoV-2 virus, not antibodies, which would reflect exposures that may have occurred weeks to months earlier [15]. The window to detect virus in infected animals is relatively short, and rRT-PCR testing outside the viral shedding window yields negative results.

SARS-CoV-2 presumably originated in an animal and spilled over into humans with subsequent spillback from humans into many other animal species. While SARS-CoV-2 remains primarily a human pathogen, known transmission to and from multiple species raises critical public health concerns and the need to identify potential amplifying or reservoir hosts, and routes of transmission. From humans, to endangered wildlife, to pets and agricultural animals, ongoing surveillance is essential as SARS-CoV-2 variants continue to emerge, creating a dynamic landscape of susceptibility and transmission risks within and between species that could have far-reaching implications on conservation, ecosystem health, and food production, in addition to economics and public health.

## Acknowledgements

This work was supported by The Ohio State University Infectious Diseases Institute and Center of Microbiome Science (Targeted Investment: eSCOUT – Environmental Surveillance for COVID-19 in Ohio: Understanding Transmission), the Centers of Excellence for Influenza Research and Response, National Institute of Allergy and Infectious Diseases, National Institutes of Health, Department of Health and Human Services under contract 75N93021C00016. We thank H. Cochran, E. Ohl, A. Cleggett, S. Treglia, A. M. Williams, F. Savona, J. W. Smith, and D. Sizemore. We are also grateful to the Ohio Department of Natural Resources, Ohio State Trappers Association, Ohio Wildlife Center, Columbus Zoo & Aquarium, Franklin County Public Health, the Ohio State University Veterinary Medical Center, MedVet Hilliard, Columbus Humane, Shelter Outreach Services of Ohio, and Ohio county fairs and metro parks for their support of this project. We further acknowledge the help of many hunters and trappers, Ohio State students, Ohio veterinarians and veterinary technicians, and community members who aided with sampling around the state of Ohio.

## Declaration of interest statement

The authors report there are no competing interests to declare.

## Author Contributions

Conceptualization: VLH, SAF, ASB

Methodology (Sample Collection and Testing): VLH, CM, DSM, JMN, DH, SL, PD, JAW, AM, RP, MF, JF, SG, RJ, ASB, VLH

Investigation (Analysis): ME, VLH, ASB, JMN

Visualization: ME

Funding acquisition: VLH, SAF, ASB

Project administration: VLH, SAF, ASB

Supervision / Consultation: VLH, PD, DD-C, ASB, JW, PY, AV, SK, QW, LJS

Writing – original draft: ME, VLH

Writing – review & editing: All

## Animal Subjects

All animal sampling conducted in this study was IACUC approved (live animal sampling) or exempt.

## Notes

### Competing Interest Statement

The authors have declared no competing interest.

